# *Chroococcidiorella tianjinensis*, gen. et sp. nov. (Trebouxiophyceae, Chlorophyta), a green alga arises from the cyanobacterium TDX16

**DOI:** 10.1101/2020.01.09.901074

**Authors:** Qing-lin Dong, Xiang-ying Xing

**Affiliations:** Department of Bioengineering, Hebei University of Technology, Tianjin, 300130, China

**Keywords:** Green algae, Cyanobacterial origin, *Chroococcidiorella tianjinensis*, 18S rRNA gene sequence, Ultrastructure

## Abstract

All algae documented so far are of unknown origin. Here, we provide a taxonomic description of the first origin-known alga TDX16-DE that arises from the *Chroococcidiopsis*-like endosymbiotic cyanobacterium TDX16 by de novo organelle biogenesis after acquiring its green algal host *Haematococcus pluvialis*’s DNA. TDX16-DE is spherical or oval, with a diameter of 2.0-3.6 µm, containing typical chlorophyte pigments of chlorophyll a, chlorophyll b and lutein and reproducing by autosporulation, whose 18S rRNA gene sequence shows the highest similarity of 99.7% to that of *Chlorella vulgaris*. However, TDX16-DE is only about half the size of *C. vulgaris* and structurally similar to *C. vulgaris* only in having a chloroplast-localized pyrenoid, but differs from *C. vulgaris* in that (1) it possesses a double-membraned cytoplasmic envelope but lacks endoplasmic reticulum and Golgi apparatus; and (2) its nucleus is enclosed by two sets of envelopes (four unit membranes). Therefore, based on these characters and the cyanobacterial origin, we describe TDX16-DE as a new genus and species, *Chroococcidiorella tianjinensis* gen. et sp. nov., which sets the basis for multidisciplinary research.

## Introduction

*Chlorella vulgaris* is the first documented *Chlorella* green alga discovered by Beijerinck in 1890 [1]. Since then, about 1000 orbicular-shaped *Chlorella* and *Chlorella*-like green algae have been isolated [2]. Classification of these small coccoid chlorophytes is a difficult task, because their origins and evolutionary histories/relationships, the crucial information necessary for their accurate assignments, are unknown. In this circumstance, the *Chlorella* and *Chlorella*-like green algae are classified traditionally by comparing the degrees of resemblance of their morphological, biochemical, physiological and ultrastructural features [3-10] and currently by inferring their origins and evolutionary histories/relationships via phylogenetic analyses of the molecular data, e.g., sequences of the SSU and ITS regions of nuclear-encoded ribosomal DNA [11-23]. As such, the taxonomy of *Chlorella* and *Chlorella*-like green algae is problematic and unstable, as indicated by the frequent taxonomic revisions.

In our previous studies, we found that the endosymbiotic cyanobacterium TDX16 resembling *Chroococcidiopsis thermalis* [24] escaped from the senescent/necrotic cells of green alga *Haematococcus pluvialis* (host) [25], and turned into a *Chlorella*-like green alga TDX16-DE [26] by de novo organelle biogenesis after hybridizing the acquired host’s DNA with its own one and expressing the hybrid genome [27]. Therefore, the *Chlorella*-like green alga TDX16-DE arises form the *Chroococcidiopsis*-like cyanobacterium TDX16, which is the first alga and also the first organism with known origin and formation process and thus of great importance and significance in biology.

The present study aims at taxonomic assignment of TDX16-DE to lay the basis for relevant research. Based upon TDX16-DE’s cyanobacterial origin in combination with its cell morphology, ultrastructure, photosynthetic pigments and 18S rRNA gene sequence, we describe TDX16-DE as a new genus and species, *Chroococcidiorella tianjinensis*, gen. et sp. nov.

## Materials and Methods

### Strain and cultivation

The green alga TDX16-DE is derived from the cyanobacterium TDX16, the formation process of which has been elucidated in our previous study [27]. The obtained TDX16-DE was inoculated into 250-ml autoclaved flasks containing 100 ml BBM medium [28] and cultivated in the incubator with continuous light of 60 μmol photons m ^−2^ &s ^−1^, at 25°C.

### Microscopy observations

Light microscopy: TDX16-DE cells were examined with a light microscope BK6000 (Optec, China). Photomicrographs were taken under the oil immersion objective lens (100×) using a DV E3 630 camera.

Transmission electron microscopy: TDX16-DE cells were harvested by centrifugation (3000 rpm, 10 min) and fixed overnight in 2.5% glutaraldehyde (50 mM phosphate buffer, pH7.2) and 1% osmium tetroxide (the same buffer) for 2 h at room temperature. After dehydration with ascending concentrations of ethanol, the fixed cells were embedded in Spurr’s resin at 60°C for 24 h. Ultrathin sections (60 to 80 nm) were cut with a diamond knife, mounted on a copper grid, double-stained with 3% uranyl acetate and 0.1% lead citrate, and examined using a JEM1010 electron microscope (JEOL, Japan).

### Pigment analyses

In vivo absorption spectrum: TDX16-DE cells were scanned with an ultraviolet-visible spectrophotometer Cary 300 (Agilent, USA).

Pigment separation and identification: Chlorophyll b (Chl b) and lutein were separated by thin-layer chromatography according to the method described by Lichtenthaler [29]. Pigments were analyzed with the spectrophotometer Cary 300 (Agilent, USA), and identified by spectroscopic matching with the published data.

### 18S rRNA gene sequence

DNA sample was prepared in the same way as described in our previous study [27]. 18S rRNA gene was amplified using the primers 5’-ACCTGGTTGATCCTGCCAGTAG-3’ and 5’-ACCTTGTTACGACTTCTCCTTCCTC-3’ [30] under the conditions: 5 min at 95°C, 35 cycles of 30 s at 95°C, 30 s at 55°C and 1min at 72°C and a final 10 min extension step at 72°C. The amplified product was sequenced on ABI 3730 DNA analyzer (PerkinElmer Biosystems, USA).

## Results

### Morphology and ultrastructure

TDX16-DE cells are green, spherical or oval, about 2.0-3.6 µm in diameter (young cells 2. 0-2.9 µm ; mature cells 2.9-3.6µm) (Fig. 1), containing one nucleus, one chloroplast, one or two mitochondria and one or several vacuoles, but no endoplasmic reticulum, Golgi apparatus and peroxysome (Fig. 2). The nucleus has two sets of envelopes, which are in close apposition in most cases (Fig. 2 B, C, D), but separated from each other when the electron-dense vesicles bud from the nuclear envelope into the interenvelope space, and migrate outside to the chloroplast envelope and cytoplasmic envelope (Fig. 2 A). The parietal chloroplast is “e”-shaped, in which there are starch granules, plastoglobules and a prominent pyrenoid that is surrounded by two starch plates and bisected by two pairs of thylakoids (Fig. 2). Mitochondria lie between the chloroplast and nucleus in the chloroplast cavity (Fig. 2 A, B, D). Vacuoles are encircled by two unit membranes, sequestering electron-transparent or electron-dense materials (Fig. 2 A, C, D). The cells are enclosed with a cytoplasmic envelope (two unit membranes), on and from which lipid droplets form and microfibrils emanate into the extracytoplasmic space respectively (Fig. 2 C, D), indicating that the double-membraned cytoplasmic envelope serves the functions of endoplasmic reticulum and Golgi apparatus for synthesizing lipids and building materials of cell wall. The cell wall comprises an inner stable trilaminar domain with a dark-light-dark configuration, reminiscent of the cell wall of TDX16 [27], and an outer loosely-compacted stratified sheath (Fig. 2), the latter of which occasionally scales off owing to osmotic-shock during cell fixation (Fig. 2C). It is noteworthy that the two unit membranes of cytoplasmic envelope and vacuoles, like those of chloroplast and mitochondrial envelopes, are difficult to distinguish when they are tightly appressed and easily misrecognized as the two layers of one unit membrane as they are separated, while the two sets of nuclear envelopes are easily misinterpreted as the two unit membranes of one nuclear envelope without being aware of their formation processes [27].

**Figure 1.**
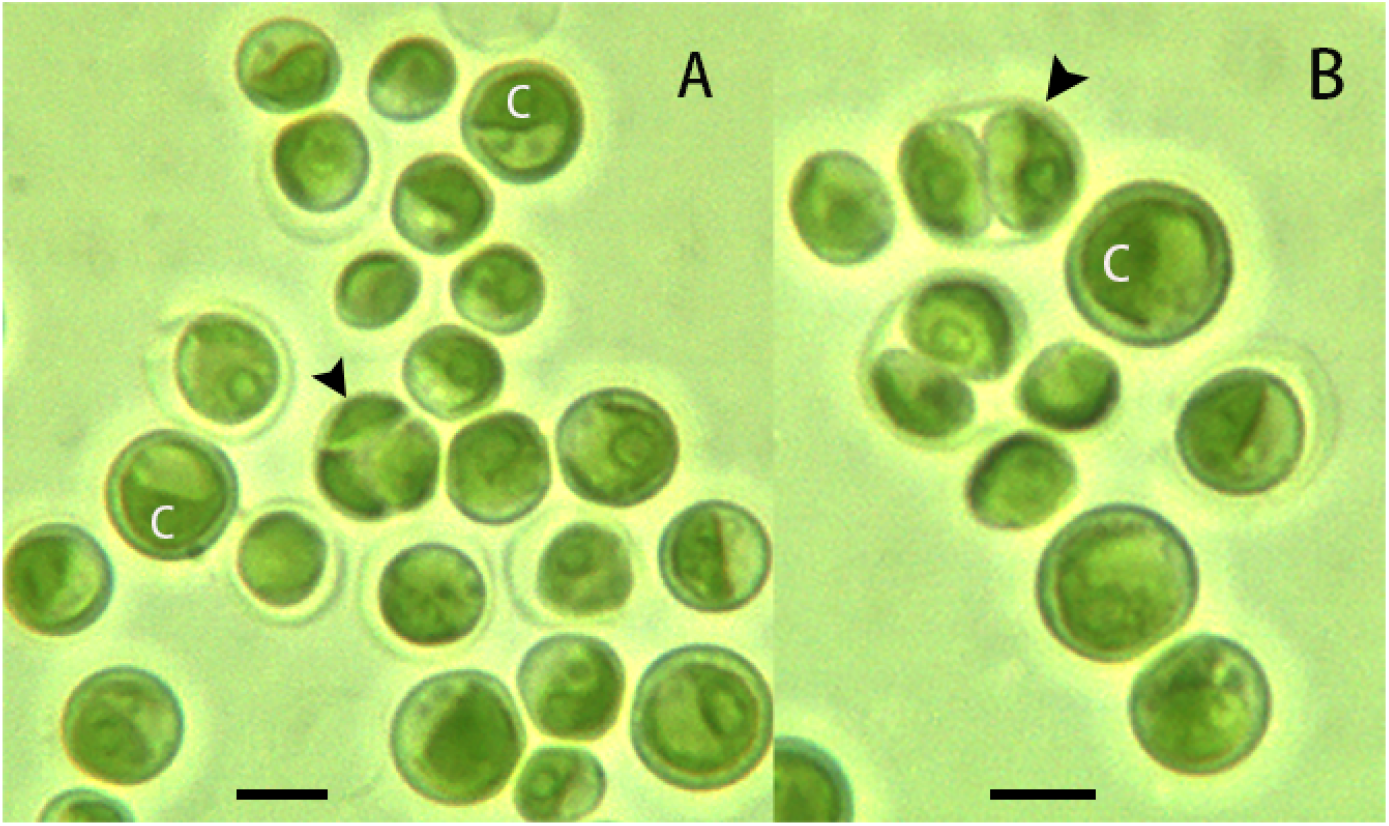
Morphology of *Chroococcidiorella tianjinensis*. (A) A three-celled autosporangium (arrowhead). (B) A two-celled autosporangium (arrowhead). C, chloroplast. Scale bars = 2 µm.

**Figure 2.**
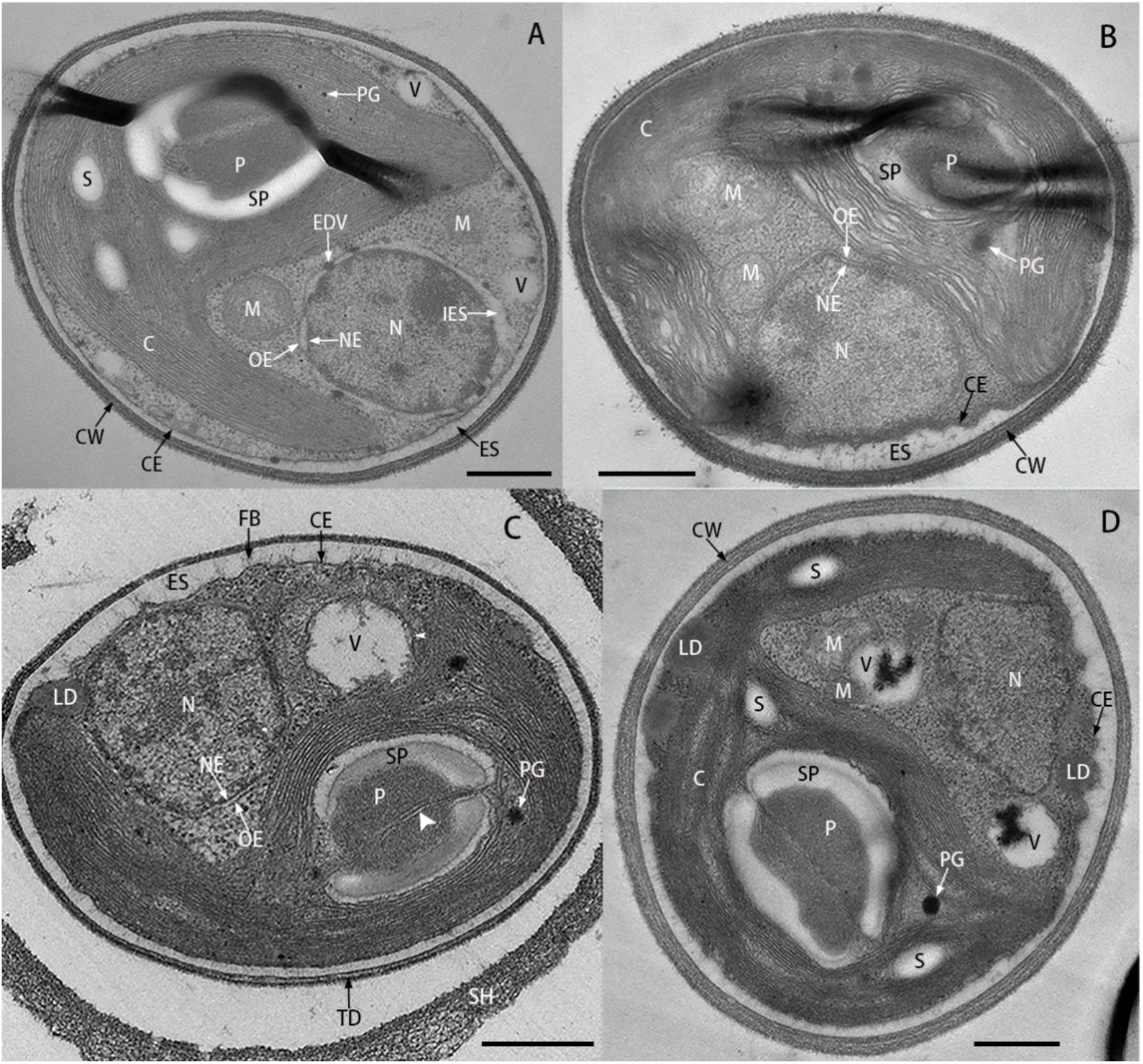
Ultrastructure of *Chroococcidiorella tianjinensis*. (A) The two sets of nuclear envelops separate from each other, resulting in a broad interenvelop space. (B) The two sets of nuclear envelopes and cytoplasmic envelope are in close apposition. (C) The pyrenoid is bisected by two thylakoids (large arrowhead), the vacuole is delimited by double membranes (small arrowhead), the outer sheath of cell wall detaches from the trilaminar domain and microfibrils emanate from the cytoplasmic envelope. (D) Lipid droplets form on the inner leaflet of cytoplasmic envelope that consists of two membranes. C, chloroplast; CE, cytoplasmic envelope; CW, cell wall; EDV, electron-dense vesicle; ES, extracytoplasmic space; FB, microfibrils; IES, interenvelope space; LD, lipid droplet; M, mitochondrion; N, nucleus; NE, nuclear envelope; OE, outer nuclear envelope; P, pyrenoid; PG, plastoglobule; S, starch granule; SH, sheath; SP; starch plate; TD, trilaminar domain; V, vacuole. Scale bars = 0.5 µm.

### Reproduction

TDX16-DE reproduces exclusively by autosporulation with the production of two (Fig. 1 B, Fig. 3 B), three (Fig. 1 A, Fig. 3 C) and four daughter cells (autospores) (Fig. 3 D) per autosporangium; the autospores are liberated after apical rupture of the autosporangium (Fig. 3 B). Structurally, the autosporangium of TDX16-DE is more similar to the sporangia of TDX16 [27] and *C. thermalis* [24] than to the aplanosporangium of *H. pluvialis* [27]. At the beginning of autosporulation, the chloroplast splits into two small ones, while the nucleus disassembles and vanishes (Fig. 3 A). Consistently, nuclei appear only in one cell in the immature four-celled autosporangium (Fig. 3 D), but in all cells in the mature three-celled autosporangium (Fig. 3 C); by contrast chloroplasts present in each cells in both the mature and immature autosporangia (Fig. 3 C, D). These results suggest that chloroplasts develop by splitting the existing ones, while nuclei form de novo in the daughter cells during autosporulation.

**Figure 3.**
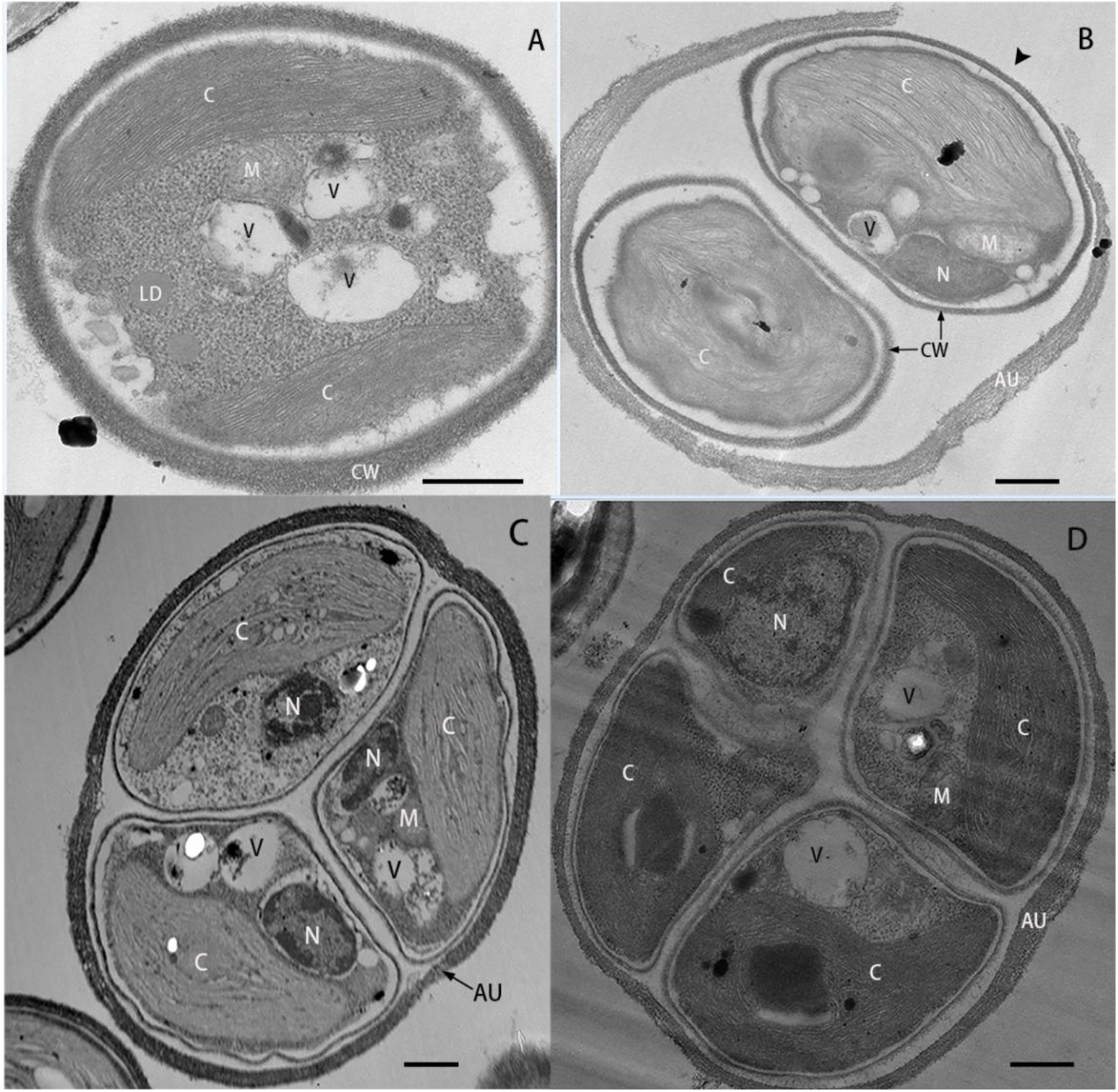
Reproduction of *Chroococcidiorella tianjinensis*. (A) A cell contains two small chloroplasts but no nucleus. (B) A two-celled autosporangium ruptures, resulting in an opening (arrowhead). (C) An autosporangium sequesters three mature daughter cells. (D) An autosporangium encloses four immature daughter cells. AU, autosporangium; C, chloroplast; CW, cell wall; LD, lipid droplet; M, mitochondrion; N, nucleus; V, vacuole. Scale bars = 0.5 µm.

### Photosynthetic pigments

In vivo absorption spectrum of TDX16-DE (Fig. 4 A) exhibits two absorption peaks of chlorophyll a (Chl a) at 440 and 680 nm, a conspicuous shoulder peak at 653 nm of chlorophyll b (Chl b) [31-32], and a merged peak of carotenoids around 485 nm. Chl b and lutein separated from TDX16-DE display absorption peaks at 456 and 645 nm (Fig. 4 B), 420, 446 and 475nm (Fig. 4 C) respectively, identical to those of the corresponding plant pigments [29]. Hence, TDX16-DE contains Chl a, Chl b and lutein.

**Figure 4.**
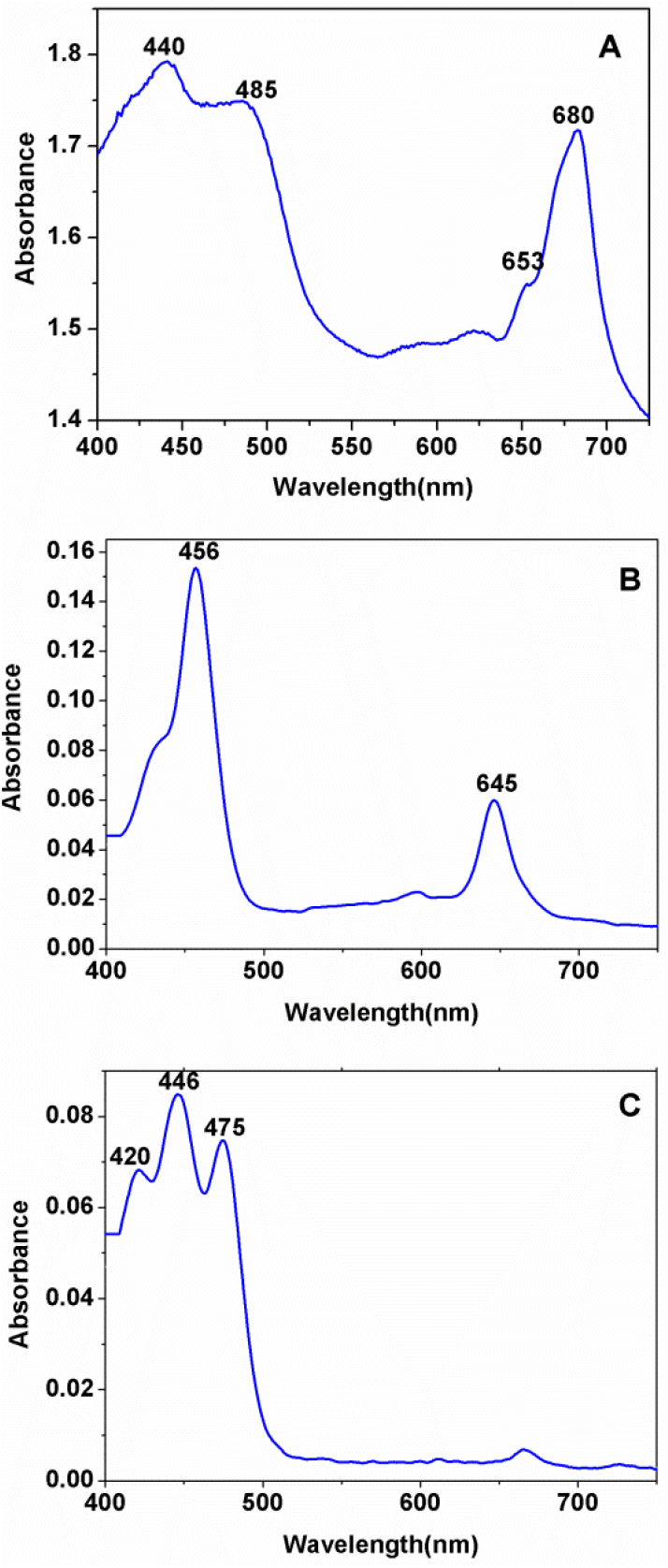
Photosynthetic pigments of *Chroococcidiorella tianjinensis*. (A) In vivo absorption spectrum. (B) Absorption spectrum of the separated Chl b. (C) Absorption spectrum of the separated lutein.

### 18S rRNA gene sequence

18S rRNA gene sequence of TDX16-DE (GenBank: MW033971.1) shows the highest similarity of 99.7% to those of *C. vulgaris* deposited in GenBank.

## Discussion

The presence of typical chlorophyte pigments of Chl a, Chl b and lutein in TDX16-DE (Fig. 4) indicates that TDX16-DE belongs to the Chlorophyta, and the highest similarity of 99.7% between the 18S rRNA gene sequences of TDX16-DE and *C. vulgaris* suggests that TDX16-DE is most similar to *C. vulgaris*. The results of cell morphology and ultrastructure, however, demonstrate that TDX16-DE is distinct from *C. vulgaris*, because they share only two common traits but differ in six characters. Their common features include (1) chloroplasts are embedded with starch granules and a pyrenoid that is surrounded by two starch plates and bisected by two pairs of thylakoids in TDX16-DE (Fig. 2) and *C. vulgaris* [33-35] and (2) reproduction is achieved by autosporulation, generating two, three and four autospores in TDX16-DE (Fig. 1, Fig. 3) and *C. vulgaris* [33, 36]. By contrast, their different characters encompass (1) TDX16-DE (2.0-3.6 µm) (Fig. 1, Fig. 2) is only about half the size of *C. vulgaris* (5.0-7.0 µm) [35-36]; (2) TDX16-DE’s chloroplast is “e”-shaped (Fig. 2); while the chloroplast of *C. vulgaris* is cup-shaped and sometimes has two lobes (bilobed) [35, 37]; (3) TDX16-DE lacks endoplasmic reticulum and Golgi apparatus (Fig. 2), while *C. vulgaris* possesses both of the two organelles [34, 37-39]; (4) TDX16-DE’s nucleus is enclosed by two sets of envelopes (Fig. 2), while the nucleus of *C. vulgaris* has one envelope [38]; (5) TDX16-DE has a double-membraned cytoplasmic envelope (Fig. 2), while *C. vulgaris* has a single-membraned cytoplasmic membrane [36]; and (6) daughter cell nuclei form de novo in TDX16-DE (Fig. 3), but develop by division of the existing one in *C. vulgaris* [36-37].

Since TDX16-DE is tiny (2.0-3.6 µm) and nearly pico-sized (< 3 µm), it is much similar in structure, particularly organelle number and arrangement, to the pico-sized *Nannochloris*-like green algae, which have a minimal set of organelles and differ from TDX16-DE in their absence of pyrenoid, such as *Choricystis minor* [40], *Pseudodictyosphaerium jurisii* [41], *Mychonastes homosphaera* [42], *Picocystis salinarum* [43], *Picochlorum oklahomensis* [44], *Chloroparva pannonica* [45] and *Pseudochloris wilhelmii* [46]. The structural similarity between TDX16-DE and these *Nannochloris*-like green algae lies primarily in three aspects: (1) absence of endoplasmic reticulum and Golgi apparatus (except the Golgi apparatus-like structure in *P. salinarum, P. jurisii* and *P. wilhelmii*); (2) the nucleus is peripherally localized in the opening of chloroplast cavity and appressed to the cytoplasmic membrane/envelope; and (3) mitochondria situate between the chloroplast and nucleus and usually attach to the chloroplast.

Phylogenetic analysis of SSU and ITS rDNA is now applied predominately for classifying the origin-unknown algae by inferring their origins and evolutionary histories/relationships, which is, however, unnecessary and also unfeasible for taxonomic assignment of TDX16-DE. Because (1) the origin and developmental relationship of TDX16-DE is clearly known; and (2) current molecular phylogenetic analyses are based on the fundamental assumption that all algae descend directly from other algae, while TDX16-DE is not the case, which arises from the cyanobacterium TDX16 by acquiring *H. pluvialis*’s DNA [27]. So, TDX16-DE descend, to some extent, only indirectly from the green alga *H. pluvialis* via the cyanobacterium TDX16. Such a developmental relationship apparently can not be inferred by current molecular phylogenetic analysis, or in other words, the inferred phylogenetic relationship does not reflect the true origin and development history of TDX16-DE (it is also the case for the common ancestor of algae that did not arise from algae).

In truth, the experimental results discussed above are sufficient for assigning TDX16-DE. The *Chlorella* and *Chlorella*-like coccoid green algae are currently categorized into two different clades within in Chlorellaceae: the *Chlorella*-clade and the *Parachlorella*-clade [13]. The highest similarity of TDX16-DE’s 18S rRNA gene sequence to that of *C. vulgaris*, and the typical *Chlorella* structure: a prominent pyrenoid surrounded by two starch plates and bisected by two pairs of thylakoids in the single parietal chloroplast, indicate that TDX16-DE falls within the *Chlorella*-clade; while TDX16-DE’s compartmental characters: two sets nuclear envelopes, double-membraned cytoplasmic envelope and vacuoles and absence of endoplasmic reticulum, Golgi apparatus and peroxysome, distinguish it from all the members within the genera of *Chlorella*-clade [2,16-17,20,23, 47]. Therefore, considering TDX16-DE’s unique compartmental features and definite origin, we propose a new genus *Chroococcidiorella* within the *Chlorella*-clade and TDX16-DE is a new species *Chroococcidiorella tianjinensis*.

### *Chroococcidiorella* Q.L. Dong & X. Y. Xing, gen. nov

Cells solitary, spherical or oval, enclosed by a double-membraned cytoplasmic envelope. Nucleus encircled by two sets of envelopes, endoplasmic reticulum and Golgi apparatus absent. One parietal chloroplast with a pyrenoid surrounded by two starch plates and traversed by two pairs of thylakoids. Asexual reproduction by autosporulation. Sexual reproduction unknown.

#### Etymology

The genus *Chroococcidiorella* is named for its origin-*Chroococcidiopsis*; plus ella, small or diminutive.

#### Type species

*Chroococcidiorella tianjinensis* Q. L. Dong & X. Y. Xing

### *Chroococcidiorella tianjinensis* Q. L. Dong & X. Y. Xing, sp. nov. (Fig. 1-3)

Cells solitary, spherical or oval, with a diameter of 2.0-3.6 µm, One nucleus with two sets of envelopes, one parietal “e”-shaped chloroplast with a pyrenoid surrounded by two starch plates and bisected by double thylakoids, double-membraned cytoplasmic envelope and vacuoles. No endoplasmic reticulum, Golgi apparatus and peroxysome. Asexual reproduction via 2-4 autospores. Sexual reproduction not observed.

#### Holotype

Strain CCTCC AF2020007^T^ has been cryopreserved in China Center for type Culture Collection, Wuhan University, Wuhan, China. Also, a living culture FACHB-3081 has been deposited in the Freshwater Algae Culture Collection at the Institute of Hydrobiology, Chinese Academy of Sciences, Wuhan, China.

#### Etymology

The species name *tianjinensis* refers to the Chinese city, Tianjin, where the formation of this alga was found.

## Conclusion

The taxonomic assignment of TDX16-DE as *Chroococcidiorella tianjinensis* sets the basis for multidisciplinary research. There is no reason to think that *C. tianjinensis* is the only alga (eukaryote) with cyanobacterial (prokaryotic) origin, so, *C. tianjinensis* along with its cyanobacterial precursor TDX16 whose genome is already available (GenBank NDGV00000000) [27] provide an unprecedented reference and platform for the study of different subjects, e.g., cell, molecular, genome, genetic and evolutionary biology, which holds the potential to resolve the relevant puzzles and make breakthroughs in biology.

